# Asynchronous trophic niche shifts among invertebrate-feeding fishes across a coral cover gradient

**DOI:** 10.64898/2026.10.15.757261

**Authors:** Zoé Delecambre, Caroline Dubé, Fabien Morat, Pauline Narvaez, Christopher P. Meyer, Emma Paul Costesec, Clément Castrec, Guillaume Iwankow, Alexandre Mercière, Jordan M. Casey, Valeriano Parravicini

## Abstract

Unraveling the links between habitat structure, resource availability and the trophic niche is key to anticipate the effects of global change on ecosystems. On coral reefs, recurrent coral bleaching and increasing anthropogenic pressures have led to rapid losses of structural complexity, with implications for food web dynamics. Invertivorous fishes represent a diverse trophic guild whose dietary niches may be particularly sensitive to variation in benthic habitat composition. Yet, whether species within this guild exhibit similar patterns of trophic niche differentiation across contrasting reef states remains poorly understood. Here, we combine otolithometry, gut content metabarcoding, and stable isotope analysis to investigate growth performance and interspecific variation in trophic niche structure among six invertivorous fish species across contrasting coral cover conditions in French Polynesia. Species exhibited differences in trophic niche breadth between high- and low-coral-cover sites, but the direction and magnitude of these differences varied among species. Despite these differences, interspecific trophic niche overlap remained largely unchanged across coral cover conditions. Rather than exhibiting synchronous trophic patterns, species within the same guild displayed contrasting niche variations across reef conditions, without consistent changes in interspecific niche overlap. These findings challenge the assumption that species sharing a trophic guild respond uniformly to habitat variation and highlight the importance of accounting for interspecific differences in trophic niches when predicting how changes in habitat structure may reorganize food webs.

## Introduction

Habitat characteristics play an important role in shaping the ecological dynamics between and within species (Nee and May 1992). Changes in habitat structure, such as fragmentation or loss of complexity, can strongly affect species distribution and resource availability. Species may adjust their resource use to these changes by broadening their diet (Colles et al. 2009) or shifting toward alternative resources (Carscadden et al. 2020). For example, terrestrial predators can broaden their diet in more heterogeneous landscapes, while coastal species such as ghost crabs may shift their resource use following habitat modifications that alter prey availability (Morrow et al. 2014). Such trophic flexibility can buffer species vulnerability to habitat degradation by allowing them to exploit alternative resources (Cloyed et al. 2021). However, the extent to which such responses are consistent across species remains unclear. As a result, the consequences of habitat degradation for community-level organization remain poorly understood.

A key uncertainty is whether trophic responses to habitat change are synchronous across species. Under the functional equivalence hypothesis (Hubbell 2005), ecologically similar species are expected to exhibit similar ecological responses, potentially leading to convergent patterns of resource use. Such synchrony could reduce niche partitioning and increase the potential for competition. In contrast, coexistence theory (Chesson 2000), emphasizes the importance of niche partitioning in maintaining species diversity and recent empirical evidence from coral reefs further highlights the importance of trophic niche partitioning for species coexistence and community biomass (Casey et al. 2026). However, whether trophic differentiation is maintained across contrasting habitat conditions, and whether species exhibit consistent or idiosyncratic patterns of resource use, remains unclear.

Coral reefs are a prominent example of ecosystems undergoing profound habitat modifications, making them an ideal system to examine the impact of habitat degradation on trophic responses. Reefs are increasingly experiencing habitat degradation and losses of structural complexity due to climate-induced coral bleaching (McClanahan et al. 2007), outbreaks of the crown-of-thorns starfish (Graham et al. 2011; Hughes et al. 2017) and overfishing (Graham et al. 2006; Eddy et al. 2021; Reverter et al. 2022; Carlot et al. 2023; Rovellini et al. 2024). These alterations can reshape benthic invertebrate communities, as coral mortality modifies habitat structure and resource availability, potentially favoring some invertebrate taxa while negatively affecting others (Alvarez-Filip et al. 2011; Taylor et al. 2018; Stella et al. 2022). These changes in benthic invertebrate communities may have bottom-up effects across coral reef food webs (Taylor et al. 2018; Brandl et al. 2025). In particular, invertivorous fishes, which rely directly on benthic invertebrates and represent one of the most diverse and abundant trophic groups on coral reefs (Delecambre et al. 2026), may experience substantial changes in resource availability and use. Yet it remains unclear whether invertivorous fishes have synchronous or species-specific responses to habitat degradation, making community-level consequences difficult to predict.

Fish diet, although strongly shaped by phylogenetic factors (Parravicini et al. 2020), is also influenced by environmental conditions. At the biogeographical level, habitat characteristics such as reef area and isolation interact with trophic niche breadth to determine species distributions across the globe (Delecambre et al. 2025). At smaller scales, habitat modification can alter feeding behavior and resource use; for example, predatory activity or foraging efficiency can decrease with habitat degradation (Schligler et al. 2021; Minier et al. 2024). Trophic responses can include niche broadening, differences in resource use or reductions in preferred prey use, with consequences for dietary overlap and competition among species (Keith et al. 2018; Semmler et al. 2021). However, these responses vary widely across species and contexts, making it unclear whether these trophic responses are synchronous across species.

In this study, we investigated growth performance and trophic niche variation of six sympatric invertivorous reef fishes across contrasting coral cover conditions in French Polynesia. Combining otolithometry (109 individuals), gut content metabarcoding (209 individuals), and stable isotope analysis (315 individuals) across eight sites on three islands, we assessed whether patterns of trophic niche breadth and interspecific overlap varied consistently among species. Due to the strong response of benthic invertebrate communities to coral mortality (Nelson et al. 2016; Sannassy Pilly et al. 2024), we expect trophic shifts to occur after disturbance. Finally, we examined whether trophic differences across habitat conditions were associated with variation in growth performance.

## Methods

### Data collection

All samples were collected in the Society Islands, French Polynesia, across sites spanning a gradient of coral cover. We sampled eight sites distributed on two high islands (Tahiti and Moorea) and one atoll (Tetiaroa), all separated by at least 1 kilometer (Fig. 1). Following the spatially heterogeneous bleaching event of 2019, Moorea was largely impacted while Tahiti remained less affected. In contrast, Tetiaroa included both impacted and less impacted sites, allowing partial disentangling of island identity and coral cover effects. Sites were classified into two broad categories of coral condition (low and high) based on visual assessments of live coral cover and associated habitat characteristics, including the extent of coral rubble and structural complexity (Graham et al. 2006, 2015). Low-condition sites were characterized by relatively low live coral cover (<10%), extensive rubble and reduced structural complexity, whereas high-condition sites exhibited greater live coral cover (∼30%) and more complex reef structures.

**Fig. 1.**
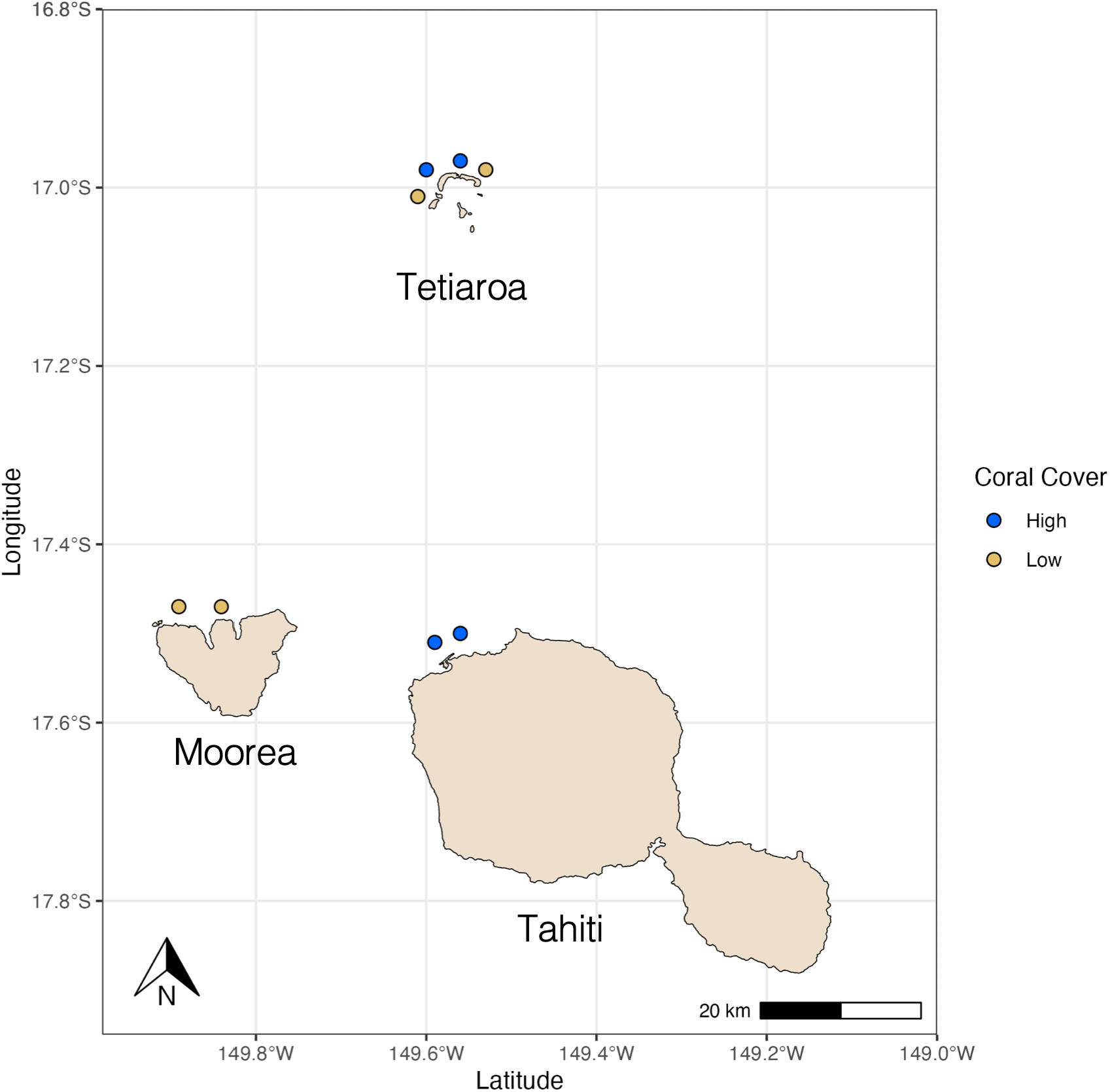
Map of the sampling sites across Moorea, Tahiti and Tetiaroa, indicating the qualified high and low coral cover sites.

At each site, we sampled 4 invertivore fish species belonging to 2 genera: *Sargocentron* (family: Holocentridae) and *Paracirrhites* (family: Cirrhitidae), with two species per genus to compare the trophic niche of closely related species. We targeted 10 individuals per species and per site, resulting in 40 individuals per site. The number of individuals sampled at each site is detailed in Table S1. In Tahiti and Moorea, we selected *Sargocentron microstoma, S. caudimaculatum, Paracirrhites fosteri* and *P. arcatus*. In Tetiaroa, *S. caudimaculatum* and *P. fosteri* were rare, so we selected *S. microstoma, S. tiere, P. arcatus* and *P. hemistictus*.

Fish were collected by spearfishing on the outer slope at 10-15m depth between 08:00 and 11:00 AM. We sampled sites around Moorea and Tahiti in April, May and June and the sites around Tetiaroa in September of 2022. Individuals were euthanized immediately after capture and stored in sealed containers filled with seawater until further processing.

### Fish dissections

Fishes were dissected under sterile conditions at the Center of Island Research and Environment Observatory (CRIOBE) field station in Moorea, in an apartment in Tahiti, and at the Tetiaroa Society research station in Tetiaroa. Each individual was photographed, weighed and measured to obtain their body mass (g) and standard length (mm). We dissected each fish to remove the entire gastrointestinal tract, then sampled stomach contents from each fish. When stomachs were empty, upper intestinal contents were sampled instead. All gut content samples were stored in sterile 2 mL tubes at −20°C, then transferred to −80°C for long-term storage until DNA extraction. White muscle tissue samples (∼1cm^3^) were taken from each individual, stored at −20°C, freeze-dried and ground for stable isotope analyses.

### DNA extraction, library preparation, and sequencing

DNA extractions were performed in the CRIOBE station in Moorea between October and November 2022. We used DNeasy PowerSoil Pro Kits (Qiagen) following the manufacturer’s protocol, with the addition two incubation steps at 65°C and a mechanical lysis of 3 cycles of 60 s at a speed of 6 m s−1 was performed with a FastPrep instrument (MP Biomedicals). DNA was quantified using a Qubit Fluoremeter (dsDNA HS Assay, Thermo Fisher Scientific, Whaltham, CA, USA). We diluted all samples to a standard concentration of 4 ng/µL. Samples with concentrations between 1 and 4 ng/µL (13.8%) were retained without dilution.

For each sample, 20 µL of DNA extract was transferred into 96 wells plates and sent to Hubbard Center for Genome Studies (HCGS) at the University of New Hampshire (UNH) for polymerase chain reaction (PCR), library preparation and sequencing. They targeted the 313 bp mitochondrial cytochrome c oxidase subunit I (COI) region using the m1COIintF and jgHCO2198 primer set (Geller et al. 2013, Leray et al. 2013), which performs well for the identification of metazoans. Following the first-round PCR amplification of the COI target region, PCR products were indexed for Illumina sequencing. For each reaction, 6.0 µL of KAPA HiFi HotStart ReadyMix was combined with 5.0 µL of dual unique index primer pairs (and 2.0 µL of diluted first-round PCR product. Indexing PCR was performed in a thermocycler with an initial denaturation at 94°C for 3 minutes, followed by 15 cycles of denaturation at 94°C for 20 seconds and extension at 72°C for 15 seconds, followed by a final extension at 72°C for 7 minutes and a hold at 4°C. The indexed PCR products were then pooled and quantified using a Qubit fluorometer. The pooled library was subjected to DNA size selection using a BluePippin system (Sage Science), following the size-selection protocol specified by the HCGS facility. The resulting size-selected pool was subsequently loaded onto an Illumina NovaSeq 6000 sequencing platform using paired end, version 1.5 chemistry on an SP flow cell. Forward and reverse read lengths were 250 base pairs. The data were demultiplexed using Illumina bcl2fastq v2.20.0.422.

### Bioinformatics

We used QIIME2 (v.2024.5, Bolyen et al. 2019) with *cutadapt* for trimming primers and filtering steps as follow: trimLeft = c(0, 0) and truncLen = c(200, 180). Paired reads were merged and chimeric sequences were removed. Ultimately, we produced an amplicon sequence variants (ASV) table. Sequence read counts ranged between 6 and 621,485 per sample. Samples with fewer than 1,000 reads were excluded from all analyses, while samples with 1,000–10,000 reads were only retained for sensitivity analyses to account for potential biases associated with inadequate sequencing depth.

Taxonomic assignment was performed by aligning sequences against the National Center for Biotechnology Information (NCBI, Sayers et al. 2026) reference database using Basic Local Alignment Search Tool (BLAST, *core_nt*, Camacho et al. 2009). Taxonomic information was retained only when percent identity reached at least 97% at the genus level and 80% at the phylum level. Any percent identity lower than 80% was designated as “unassigned”.

We removed all singleton ASVs (i.e., ASVs with only one sequence read across the entire dataset). To minimize non-target amplification and potential contamination, we removed all ASVs with taxonomic assignments to the families of the host fish consumers (i.e. Holocentridae and Cirrhitidae) and terrestrial mammals.

### Stable isotopes analysis

White muscle tissue samples were freeze-dried for 48 hours at the CRIOBE station in Moorea. Dried samples were ground using two metal beads in a TissueLyser (Qiagen). Approximately 1 mg of homogenized material per sample was analyzed at the Cornell University Stable Isotope Laboratory (COIL) using a Thermo Delta V isotope ratio mass spectrometer (IRMS) coupled to a NC2500 elemental analyzer (May-June 2024). Stable isotopes ratios (‰) of δ^15^N, 8^13^C and 8^34^S were obtained for 315 individuals across the six species. δ^15^N provides information on trophic position, while δ^13^C reflects the relative contribution of pelagic *versus* benthic carbon sources. δ^34^S was used to further characterize resource origin and habitat use.

We focused on within-site comparisons, so we did not collect baseline isotope values from end members. We therefore did not compare raw isotopic data across sites or islands. Because individuals within each site share a common isotopic baseline, this approach allows robust comparisons of niche position and overlap among species. We combined δ^15^N, δ^13^C and δ^34^S values to characterize the position and extent of isotopic niches within each site.

### Otolithometry

Otoliths were used to estimate growth, as a proxy of individual performance. Sagittal otoliths were extracted during dissection, rinsed, dried with paper towels, and stored in 2 mL tubes.

At the CRIOBE in Perpignan, France, otoliths were included in small silicone molds with 20/20 epoxy resin. After 24 hours of solidification, otoliths were sliced around the core with a T210 precision crosscut to obtain a 1-millimeter slice. The slices were then polished and read under a microscope to estimate precisely the number of growth marks and the distance between growth marks and the core.

### Data analysis

A summary of the number of samples analyzed for each data type (metabarcoding, stable isotopes, and otoliths) is provided in Table S1. As a result of the 2019 bleaching event, coral cover and island identity are confounded in Moorea and Tahiti. Therefore, inferences on coral cover effects cannot be separated from island effects in these sites. In contrast, Tetiaroa includes both high and low coral cover sites, allowing evaluation of coral cover effects independently of island identity. Analyses were therefore conducted separately for Tetiaroa and the Moorea-Tahiti dataset.

#### i) Plasticity

To quantify trophic plasticity in the invertivorous fishes, we combined information on diet (gut content metabarcoding and stable isotopes) with estimates of individual growth performance. Changes in diet were interpreted alongside estimates of growth performance, allowing us to evaluate whether trophic differences co-occurred with changes in individual performance.

##### a. Multivariate analyses - Plasticity

Gut content metabarcoding data were primarily analyzed with presence-absence information to characterize resources use, while relative read abundance (RRA) was used to compare the relative contribution of prey items across samples. To explore the general patterns of trophic variation between species and coral cover conditions, we performed a principal coordinate analysis (PCoA) with the R package *ape* (Paradis and Schliep 2019) on the gut content metabarcoding dataset. We then used multivariate generalized linear models (*manyglm* function in the R package *mvabund*; Wang et al. 2022) to test whether trophic composition varied with coral cover, species identity, and their interaction. The interaction term allowed us to assess whether species differed in their trophic response to habitat condition. Separate models were fitted for gut content metabarcoding and stable isotope data.

##### b. Growth rate and coral cover

Growth information was used as input to estimate growth with the *fishgrowbot* R package (Schiettekatte 2021). We first used the *bcalc* function to perform the back-calculations and obtain length-at-age estimations, and then we used the *growthreg* function to fit the von Bertalanffy growth model. For each individual of each species, we estimated K, the rate at which the studied individual approaches the asymptotic size in its population and L∞, the theoretical maximal size in the population (cm).

#### ii) Synchrony

To quantify the degree of synchrony in trophic responses among sympatric species, we evaluated how trophic niche breadth and interspecific niche overlap varied with coral cover. Synchronous responses are expected to result in similar changes in niche breadth and higher overlap among species, whereas species-specific responses should lead to divergent patterns and limited changes in overlap. We compared gut content metabarcoding and stable isotope data to obtain complementary estimates of trophic niche structure.

##### a. Trophic niche breadth and overlap with gut content metabarcoding

Trophic niche breadth was estimated at the individual level as ASV richness and as the phylogenetic diversity of consumed resources. A phylogenetic tree of prey ASVs was built using the function *pd* from the R package *picante* (Kembel et al. 2010). To explore the effect of coral cover on niche breadth at the species level, we then performed Bayesian mixed-effect models for each species, with phylogenetic diversity and richness as the response variables, coral cover as the explanatory variable, and site as a random effect. Bayesian models were implemented with the R package *brms* (Bürkner 2021), using default priors, 2000 iterations and 4 chains, removing the first 1000 iterations as burn-in. For each model, we examined all posterior predictive distributions and trace plots.

Trophic niche overlap between pairs of individuals was estimated based on the proportion of shared prey items (presence–absence data) between individuals of different species. To evaluate how coral cover influenced interspecific overlap, we fitted Bayesian mixed-effect models for each species, with niche overlap as the response variable, coral cover as the predictor, and site as a random effect.

##### b. Trophic niche bredth and overlap with stable isotopes

Stable isotopes ratios (δ^15^N, 8^13^C and 8^34^S) were used to define the region and breadth of the trophic niche. Because isotopic baselines may vary among sites, all analyses were conducted at the site level. We used the R package *nicheROVER* (Lysy et al. 2023) to define the breadth of the trophic niche of the 4 sympatric species at each site. Interspecific niche similarity was estimated as the overlap between isotopic niches of two sympatric species within the same site with the *maxLikOverlap* function in the R package *SIBER* (Jackson and Parnell 2023).We performed two sets of Bayesian linear models for each species with the R package *brms* (Bürkner 2021): one set with niche breadth as the response variable, and one set with the overlap as the response variable. Both sets had coral cover as the explanatory variable. Again, we used 2000 iterations and 4 chains, removing the first 1000 iterations as burn-in.

## Results

The final metabarcoding dataset included 22,324,503 sequence reads and 1,369 ASVs, spanning 20 phyla, across 209 samples (Fig. S1A). On average, the sequencing depth was 104,320 reads and ASV diversity was 61.42 ASVs per individual. After excluding the unassigned ASVs (30.3%), diet composition was dominated by Arthropoda (37.2%), Chordata (34.5%) and Apicomplexa (16.7%), whereas all other phyla were rare (<1%) across all fish species. Diet diversity varied among individuals, ranging from a single phylum (18.1% of individuals) to eight phyla consumed by a *S. microstoma* individual in Tetiaroa.

Within sites, trophic niches were largely partitioned among species (Fig. S1B). At fine taxonomic resolution, most resources were not shared among species (on average, 70.8% of genera, 71.2% of species and 87.9% of ASVs were unique to a given species). Even at the phylum level, resource overlap remained limited, with 35.2% of phyla not shared among species and only 19.2% shared by all four co-occurring species.

### i) Plasticity

Broad dietary changes were explained by all three tested predictors: consumer species, coral cover (and/or island in ‘Moorea-Tahiti’ models) and the interaction between consumer species and coral cover, except for the stable isotopes model in Tetiaroa (Table T1, Fig. S2). Regardless of consumer species, ASV richness did not markedly change between high and low coral cover sites, although we detected a slight increase in low coral cover sites (high = 851 unique ASVs; low = 897 unique ASVs).

**Table T1.** Model outputs for each method (DNA metabarcoding *versus* stable isotope analysis) and island (Tetiaroa *versus* Moorea-Tahiti). Bold p-values indicate significant effects.

| Data | Island | Predictor | Degrees of freedom | Deviance | P-value |
| --- | --- | --- | --- | --- | --- |
| **Gut metabarcoding** | **Tetiaroa** | Species | 98 | 2144.50 | **0.023** |
|  |  | Coral cover | 97 | 1502.70 | **0.001** |
|  |  | Species × coral cover | 94 | 317.20 | **0.001** |
|  | **Moorea-Tahiti** | Species | 103 | 3202.00 | **0.002** |
|  |  | Coral cover | 102 | 1560.00 | **0.001** |
|  |  | Species × coral cover | 99 | 584.00 | **0.001** |
| **Stable isotopes** | **Tetiaroa** | Species | 158 | 253.12 | **0.002** |
|  |  | Coral cover | 157 | 39.13 | **0.002** |
|  |  | Species × coral cover | 154 | 14.34 | 0.146 |
|  | **Moorea-Tahiti** | Species | 146 | 241.03 | **0.002** |
|  |  | Coral cover | 145 | 143.04 | **0.002** |
|  |  | Species × coral cover | 142 | 31.92 | **0.002** |

We estimated growth performance for 30 individuals of *S. microstoma*, 20 individuals of *S. caudimaculatum*, 20 individuals of *S. tiere*, 16 individuals of *P. arcatus*, 13 individuals of *P. fosteri* and 10 individuals of *P. hemistictus*. Overall, habitat condition (coral cover) rarely impacted growth. In *P. fosteri*, most individuals from the high coral cover sites showed lower growth performance compared to individuals from the low coral cover sites (Fig. 2A). Maximum individual size (L∞) showed no difference between high and low coral cover sites across most species, except for *P. fosteri* where maximum body sizes were lower in individuals sampled in low coral cover sites (Fig 2B).

**Fig. 2.**
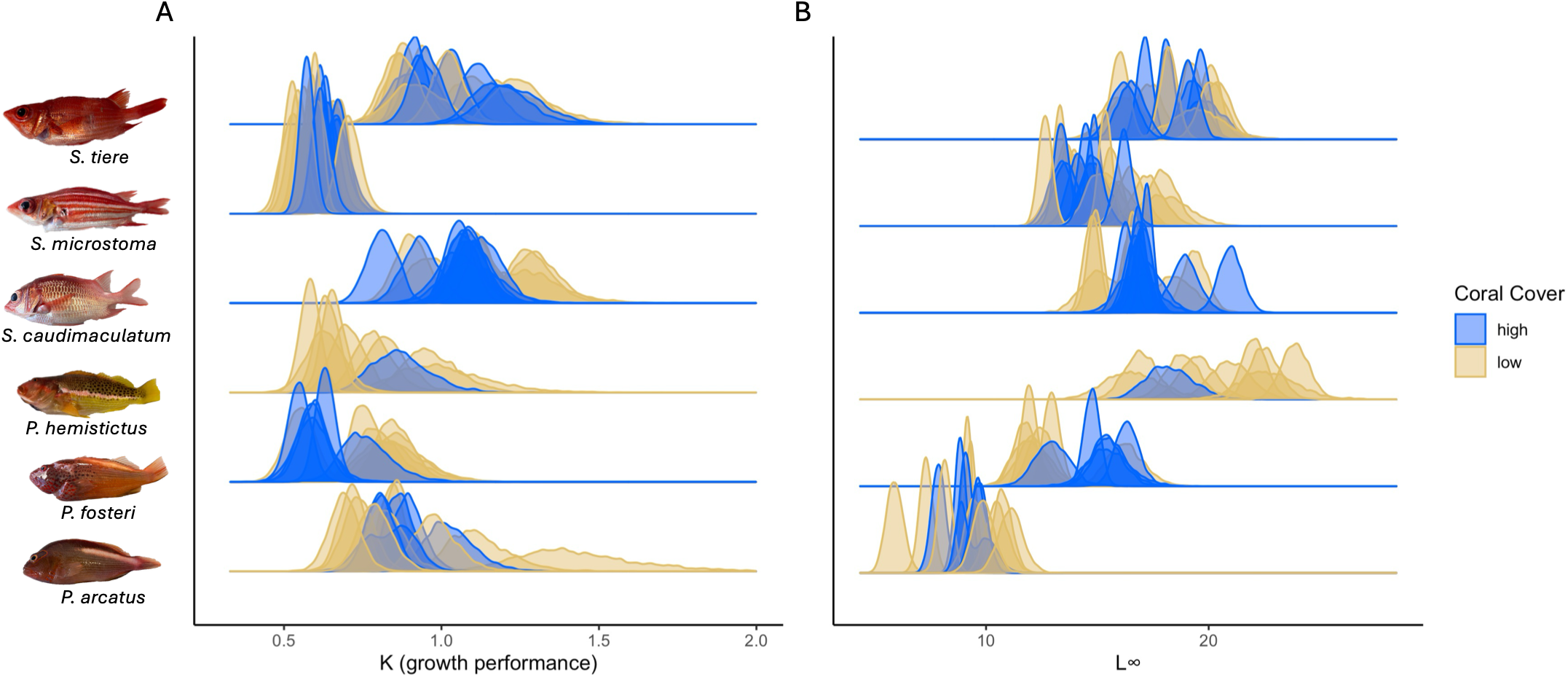
Growth model posterior distributions of (A) K and (B) L∞ of six invertivorous fishes at high and low coral cover sites. K represents the estimated rate at which the individual approaches the asymptotic size in its population, and L∞ represents the theoretical maximum body size in the population (cm). In both plots, each density curve represents an individual.

### ii) Synchrony

Trophic niche breadth showed high variability among individuals, with phylogenetic diversity ranging from 0 to 9.9 (*Paracirrhites arcatus* from a low coral cover site in Moorea) and ASV richness up to 83 distinct ASVs (*P. arcatus* from a low coral cover site in Tetiaroa). Across individuals, mean phylogenetic diversity was 1.99, and mean ASV richness was 15.

In Moorea and Tahiti, species exhibited contrasting responses to coral cover, with no common pattern across taxa. *P. arcatus* tended to increase its niche breadth with lower coral cover, while *P. fosteri* showed the opposite trend. Only *S. caudimaculatum* and *S. microstoma* displayed a similar reduction in niche breadth (Fig. 3A). In Tetiaroa, no niche breadth variation was observed in response to coral cover. However, both species increased niche breadth with lower coral cover in *Sargocentron*, while both *Paracirrhites* species had higher phylogenetic diversity at low coral cover sites (Fig. 3C). While the trophic niche of *P. arcatus* was similar between Moorea-Tahiti and Tetiaroa, the niche breadth of *S. microstoma* and *P. fosteri* in Moorea-Tahiti was higher at low coral cover sites, although only the increase observed in *S. microstoma* was notable (Fig. 3B). In Tetiaroa, no changes were detected, and trends were weak and species-specific (Fig. 3D).

**Fig. 3.**
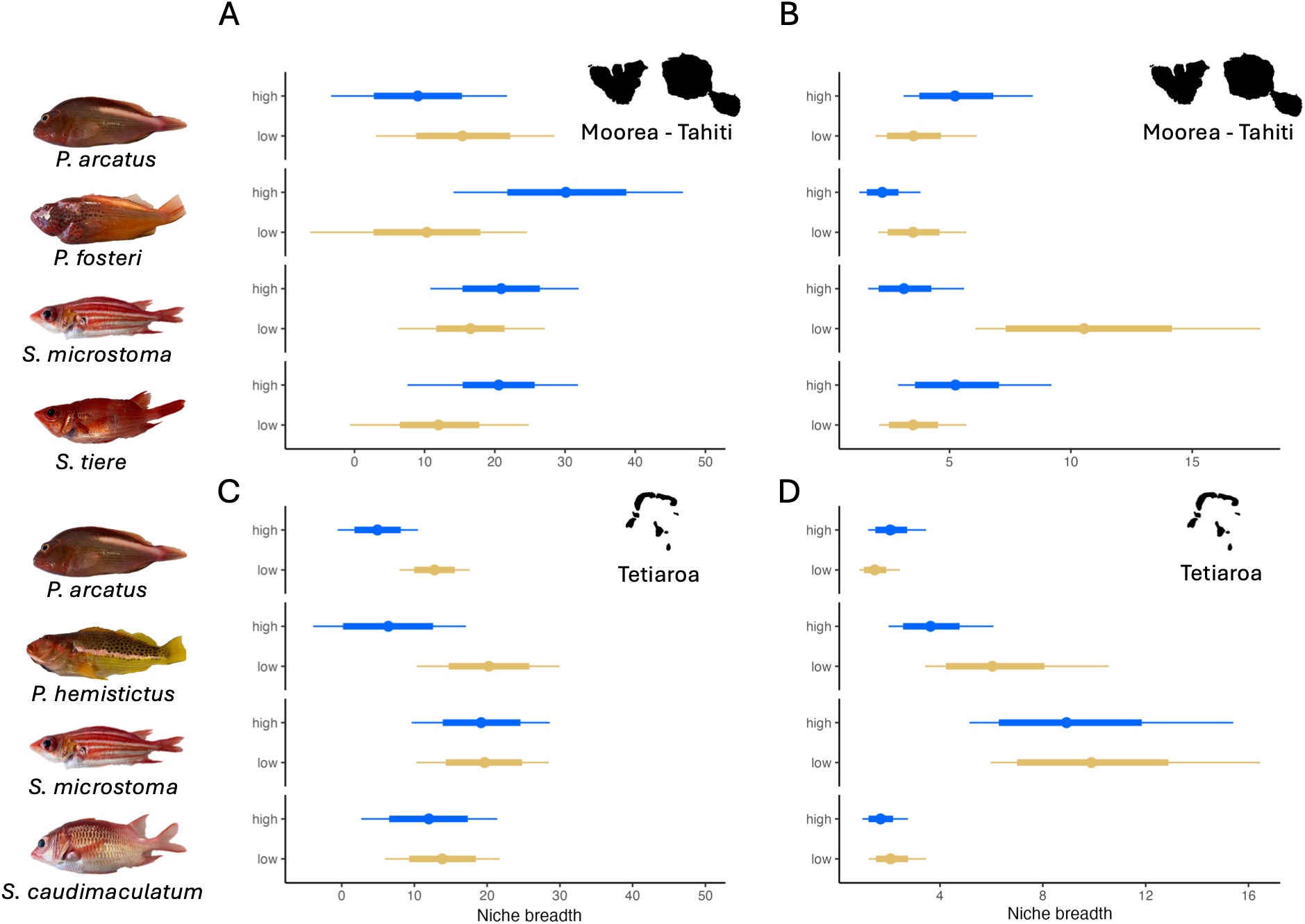
The trophic niche breadth of four invertivorous fish species in (A-B) Moorea-Tahiti and (C-D) Tetiaroa, demonstrating the (A, C) phylogenetic diversity of resources from dietary metabarcoding (B, D) and isotopic niche breadth in individuals from high and low coral cover sites. Dots represent the mean, thick lines represent 75% of the posterior distribution, and thin lines represent 95% of the posterior distribution.

Across all species, estimates of interspecific trophic niche overlap based on gut content metabarcoding showed no difference between high and low coral cover sites. In Moorea-Tahiti, responses were weak and species-specific, with *Sargocentron caudimaculatum* showing slightly higher overlap and *Paracirrhites arcatus* lower overlap, although these trends were not remarkable (Fig. 4A). In Tetiaroa, only *P. hemistictus* showed a higher overlap with low coral cover (Fig. 4C), while no change was observed for *S. microstoma* across sites.

**Fig. 4.**
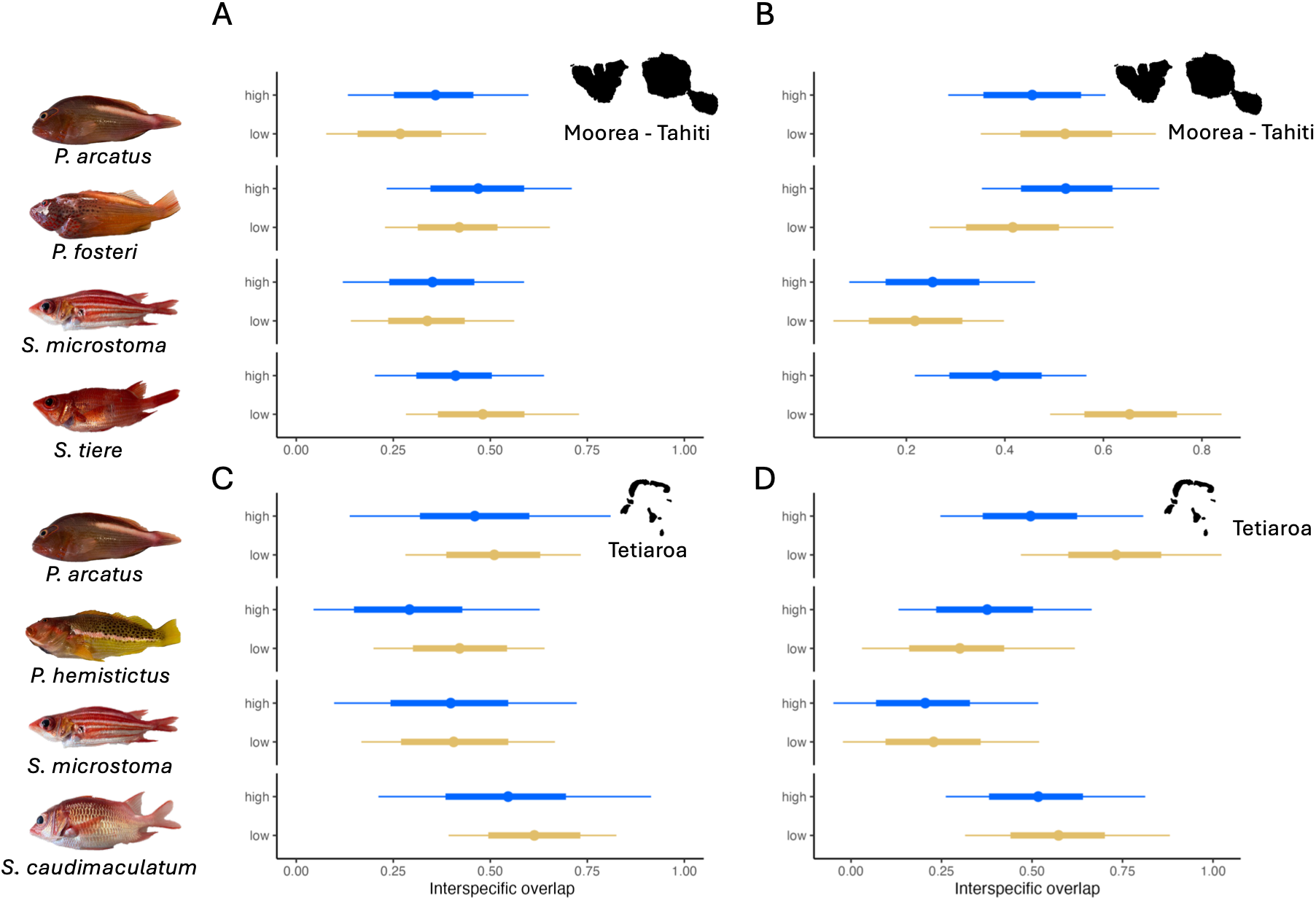
The competition model outputs for four invertivorous species in (A-B) Moorea-Tahiti and (C-D) Tetiaroa), showing (A, C) dietary metabarcoding (B, D) and isotopic trophic niche overlap in individuals from high and low coral cover sites. Dots represent the mean, thick lines represent 75% of the posterior distribution and thin lines represent 95% of the posterior distribution.

Isotopic niche overlap patterns mirrored these results. In Moorea-Tahiti, only *S. caudimaculatum* showed slightly higher in niche overlap (Fig. 4B), while in Tetiaroa, slightly higher overlap was observed for *P. arcatus* (Fig. 4D). Overall, changes in interspecific overlap were limited and inconsistent across species.

## Discussion

Combining gut content metabarcoding and stable isotope analysis, we found that the trophic niches of six invertebrate-feeding fishes differed between high and low coral cover sites in French Polynesia, although the direction and magnitude of these differences varied among species. Analyses of niche breadth and interspecific overlap revealed no consistent patterns across the invertivore guild. Instead, species exhibited idiosyncratic, and sometimes population-specific, variation in their trophic niches. Despite these differences, growth performance showed limited variation across habitat conditions, suggesting that variation in trophic niches was not associated with substantial differences in growth. Together, these results indicate that trophic responses to habitat condition may be heterogeneous across species, despite occurring within the same trophic guild.

Habitat degradation alters communities of benthic invertebrates, leading to substantial turnover in species composition (Sánchez-Fernández et al. 2012; Clever et al. 2024). Degraded coral structures can support high densities and diversity of mobile invertebrates (Nelson et al. 2016; Levy et al. 2024). Across the Caribbean, Red Sea, and Indo-Pacific, dead corals systematically support the highest mobile invertebrate density and taxonomic richness compared to living colonies regardless of overarching reef health (Caribbean: Nelson et al. 2016; Dubé et al. 2023; Red Sea; Levy et al. 2024; Indo-Pacific: Stella et al. 2022; Kamen et al. 2024). The remaining of structural complexity in dead colonies and epibiont colonization on the surface of dead corals may create habitat, food resources and settlement cues for invertebrate communities (Nelson et al. 2016). Our survey does not directly explore the community composition of invertebrates within coral colonies; however, estimating invertebrate diversity through gut content analysis of invertebrate-feeding consumers is a potential approach to approximate invertebrate diversity (e.g., Kamen et al. 2024).

The large diversity of resources (here, invertebrates) hosted by degraded coral colonies and coral rubble could also explain the high variability of trophic responses in fish consumers. Higher inter- and intra-specific niche variations have been observed in environments with higher prey diversity (Sanchez-Hernandez et al. 2020). Indeed, rather than a consequence, inter and intra-specific competition could be a cause for niche partitioning among or between species, especially with various resources availability, promoting trophic specialization (Araújo et al. 2011; Ferreira et al. 2025). However, competition and prey diversity alone cannot solely predict the response of consumers. Predation pressure, also known as top-down control, can influence the extent and the direction of niche shifts under changing environmental conditions (Sánchez-Fernández et al. 2012; Jones and Post 2016; Ferreira et al. 2025). As such, species experiencing high predation pressure tend to constrict their trophic niche in the face of change, while species with low predation pressure broaden their trophic niche (Jones and Post 2016).

The combination of differences in trophic niches across habitat conditions and largely unchanged growth performance is consistent with the possibility that trophic flexibility allows these species to exploit different resources without substantial costs to growth. Observed changes ranged from changes in resource use to trophic niche expansion, suggesting that invertebrate-feeding species can adjust their trophic strategy to exploit alternative resources under habitat degradation. While some highly specialized reef fishes show no evidence of trophic plasticity (e.g. obligate corallivores; Semmler et al. 2025) trophic plasticity has been reported in other reef fish species, as well as across trophic guilds, including invertivores, omnivores, corallivores and top predators (Lawton et al. 2012; Liedke et al. 2016; Matich et al. 2017; Clever et al. 2024). While trophic plasticity may be beneficial to maintain individual performance (Davis et al. 2022), it may also involve trade-offs, such as increased foraging cost or reduced assimilation efficiency (Hauss et al. 2023; Liu et al. 2025). However, the limited variation in growth performance suggests that these constraints are not strong in our time-for-space study system.

The contrasting patterns of trophic niche variation among invertivorous fishes, together with the absence of a systematic increase in interspecific niche overlap, suggest that trophic differentiation may be maintained across contrasting reef conditions, potentially contributing to the persistence of trophic functions under habitat degradation. Species-specific responses to environmental variations have been extensively observed in corals (Okazaki et al. 2017) and trees (Klanderud 2008; Fei et al. 2017). In vertebrates, species-specific changes have been shown in birds (Pacifici et al. 2017), bats (Reusch et al. 2019) and other mammals (Pacifici et al. 2017). In reef fishes, idiosyncratic trophic variation has been observed in two reef fish species, *Chaetodon capistratus* and *Hypoplectrus puella*, across a gradient of coral cover in the Caribbean (Clever et al. 2024). Thus, rather than exhibiting uniform trophic patterns across habitat conditions, species may differ in their resource use, potentially contributing to the maintenance of trophic niche partitioning and, ultimately, species coexistence. Indeed, the divergence of trophic responses between species in the face of the same external condition limits interspecific overlap and competition, with species shifting away from targeting the same desirable but limited resources (Ferreiro-Arias et al. 2021; Cong et al. 2023). This phenomenon decreases the risk of competitive exclusion, where outcompeted species tend to relocate or go extinct, hence promoting coexistence even between sympatric species (Day et al. 2016; Cong et al. 2023).

Our sampling included only six species, representing a subset of the invertivore guild. Extending this approach to a broader range of taxa would strengthen the generality of our findings. While focusing on invertivorous fishes is particularly relevant given the sensitivity of benthic invertebrate communities to changes in coral habitats, other trophic guilds (e.g. herbivorous or detritivorous fishes) may exhibit different patterns across habitat conditions. Additionally, our estimates of growth performance and interspecific trophic interactions rely on indirect measures. Otolith-derived growth estimates provide information on individual growth histories but do not capture other components of performance, such as survival or reproductive success. Similarly, trophic niche overlap reflects shared resource use but does not directly quantify competition, which also depends on resource availability. Although these approaches are widely used in ecological studies (Swanson et al. 2015; Martino et al. 2019; Larocque et al. 2021; Xieu et al. 2021) direct observations and experimental approaches would help clarify the mechanisms underlying the observed patterns.

## Conclusion

By combining gut content metabarcoding, stable isotope analysis, otolithometry, and coral cover, we offer a multifaceted perspective on fish responses to habitat change. As coral reefs continue to face global decline, integrative approaches are essential to understand how fish communities reorganize and persist. Our findings underscore the need to move beyond single-species or single-metric assessments and highlight the importance of integrating species-specific trophic variation into guild-level approaches to understand how coral reef food webs may reorganise under habitat degradation. Importantly, the observed responses suggest that habitat degradation may not uniformly translate across species or trophic groups, but rather reshape ecological interactions and resource use. This highlights the importance of considering changes in feeding strategies and trophic pathways alongside changes in coral cover when assessing the ecological consequences of reef degradation. Such complementary indicators may also provide earlier signals of functional change than structural habitat metrics alone, improving our ability to detect emerging shifts in ecosystem functioning. Our results support a shift toward assessing reef resilience not simply as the persistence of species, but as the ability for ecological functions and interactions to be maintained or reorganized under increasing environmental stress.

## Supplementary material

**Table S1.** Summary of the three sample types (metabarcoding, stable isotopes, and otoliths) used in the analyses, sorted by island and coral cover. DNA metabarcoding only includes samples with greater than 10,000 reads.

|  | **Moorea** | **Tahiti** | **Tetiaroa** | **High coral cover** | **Low coral cover** | **Total** |
| --- | --- | --- | --- | --- | --- | --- |
| DNA Metabarcoding | 54 | 53 | 102 | 99 | 110 | 209 |
| Stable Isotopes | 78 | 74 | 163 | 155 | 160 | 315 |
| Otoliths | 33 | 30 | 46 | 43 | 66 | 109 |

**Fig. S1.**
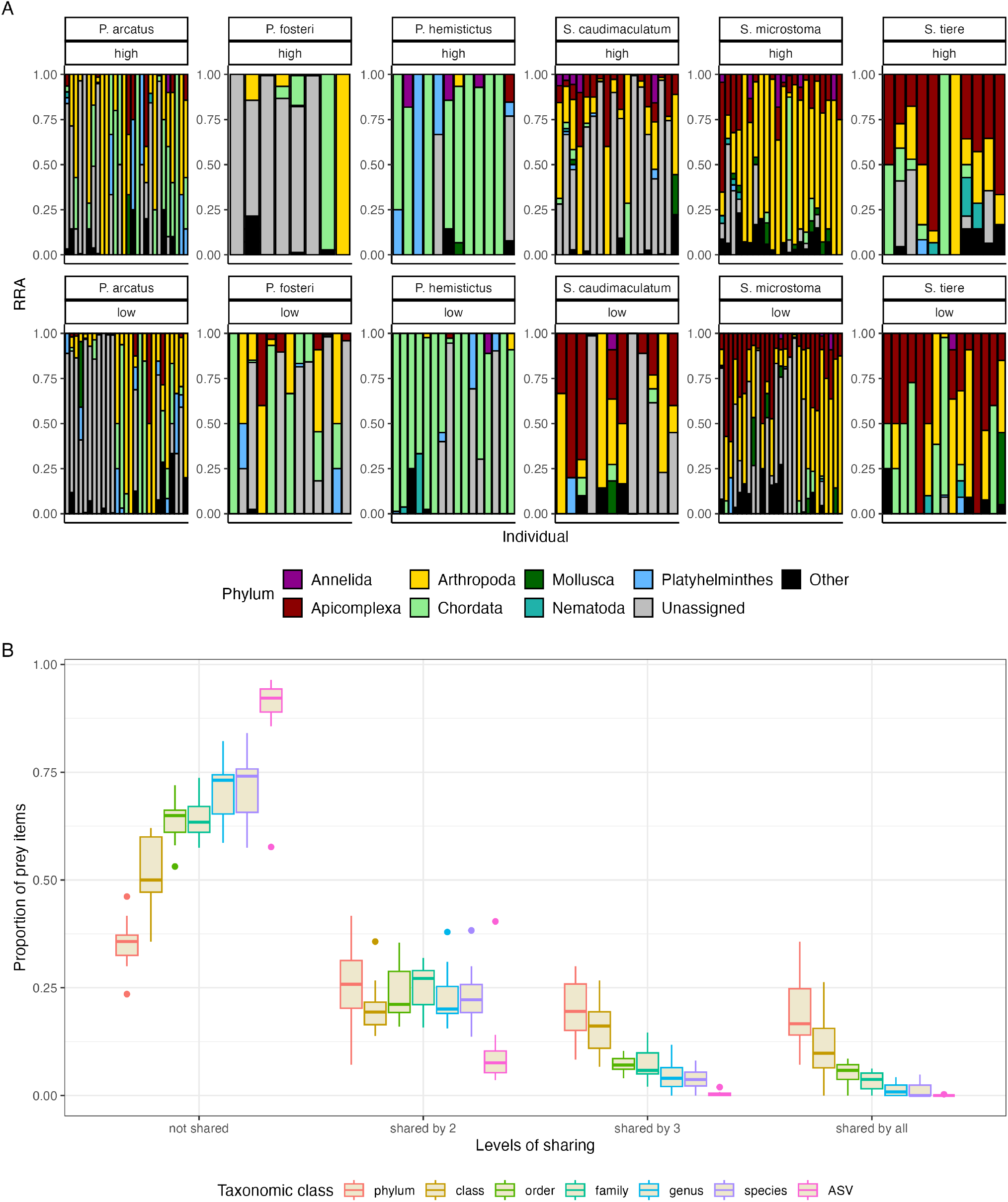
(A) Relative read abundance of prey phyla for the six invertivorous fish species in high (top) and low (bottom) coral cover sites. (B) Proportion of unshared and shared prey items at each taxonomic level (phylum to species), and at the ASV-level, categorized by whether a prey item is unique to one species or shared by two, three, or four species co-occurring at a site.

**Fig. S2.**
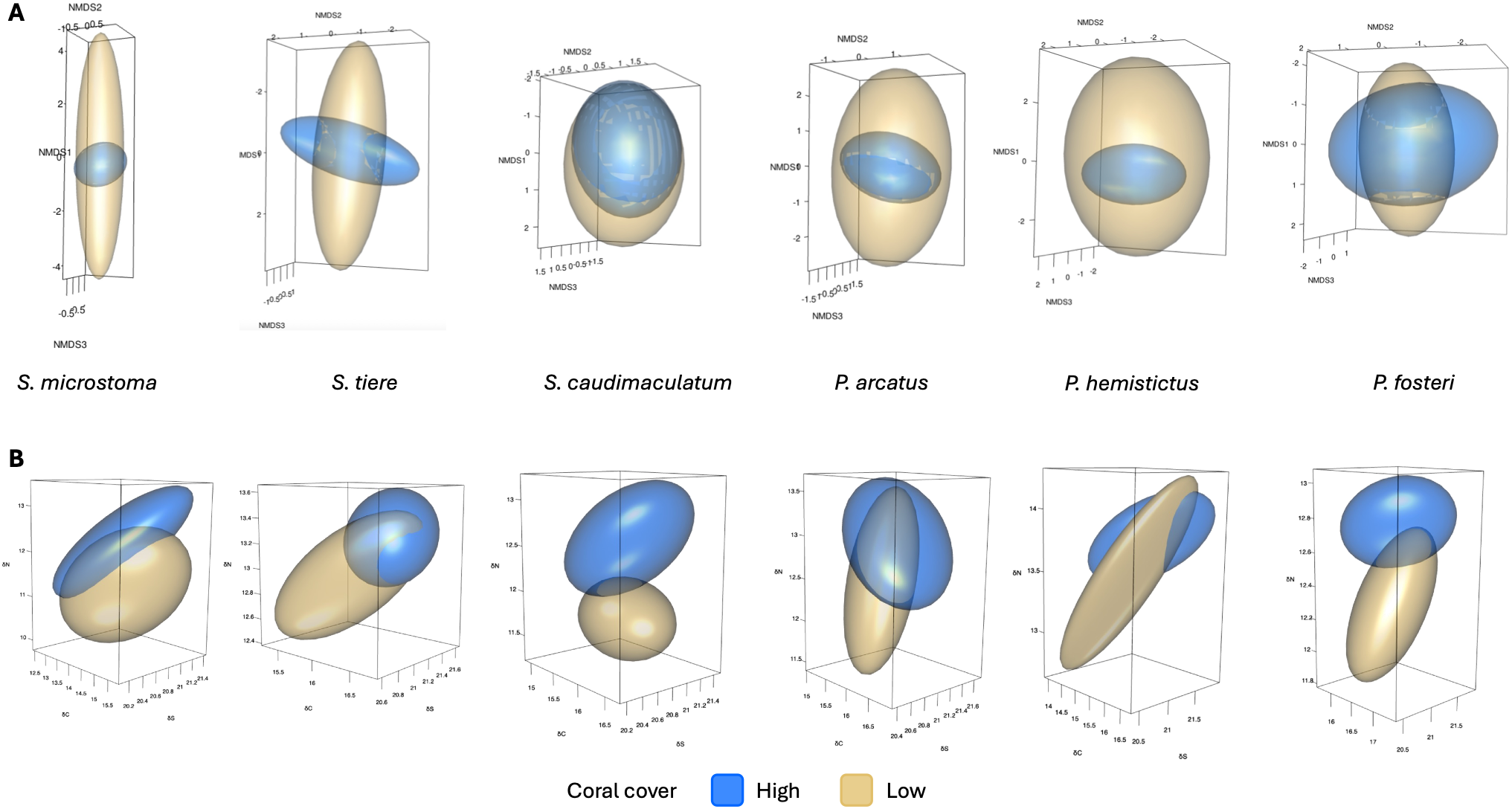
Trophic niche volumes of six invertivorous fish species across all sites based on (A) gut content metabarcoding and (B) stables isotopes.

